# Efficient Ribosomal Rna Depletion from *Drosophila* Total Rna for Next-Generation Sequencing Applications

**DOI:** 10.1101/2024.11.28.625868

**Authors:** Omkar Koppaka, Shweta Tandon, Ankita Chodankar, Awadhesh Pandit, Baskar Bakthavachalu

## Abstract

We developed a cost-effective enzyme-based rRNA-depletion method tailored for *Drosophila melanogaster*, addressing the limitations of existing commercial kits and the lack of peer-reviewed alternatives. Our method employs single-stranded DNA probes complementary to *Drosophila* rRNA, forming DNA-RNA hybrids. These hybrids are then degraded using the RNase H enzyme, effectively removing rRNA and enriching all non-ribosomal RNAs, including mRNA, lncRNA and small RNA. When compared to a commercial rRNA removal kit, our approach demonstrated superior rRNA removal efficiency and mapping percentage, confirming its effectiveness. Additionally, our method successfully enriched the non-coding transcriptome, making it a valuable tool for studying ncRNA in *Drosophila*. The probe sequences and rRNA-depletion protocol are made freely available, offering a reliable alternative for rRNA-depletion experiments.

## INTRODUCTION

*Drosophila melanogaster* has emerged as a key model organism due to rapid generation time, easy maintenance, well-characterized genome and low experimental cost (Mirzoyan et al., 2019; Tolwinski, 2017). With a genome that shares ∼60% homology with humans of which 75% are linked to human diseases (Adams et al., 2000; Banfi et al., 1996; Fortini et al., 2000), *Drosophila* plays a critical role in studying numerous conditions such as neurodegeneration, cardiac diseases, metabolic disorders and even rare human diseases (Goodman & Bellen, 2022; Kim et al., 2021; Link & Bellen, 2020; Souidi & Jagla, 2021). Beyond disease models, *Drosophila* is also essential for exploring normal physiology and gene regulation, as its transcriptome provides insights into conserved cellular mechanisms and gene interactions fundamental to development and cellular function (Hardin et al., 1990; Harvey et al., 2003; Nüsslein-Volhard & Wieschaus, 1980). This model organism enables researchers to investigate gene regulation under a range of conditions, advancing our understanding of both healthy and pathological states.

RNA expression plays a crucial role in maintenance of physiological balance, as evidenced by the fact that dysregulation of RNA leads to several pathological conditions (Bradley & Anczuków, 2023; Nemeth et al., 2024). To understand the roles of RNA expression in physiological and pathological conditions, identifying both global and tissue-specific gene expression patterns is crucial. Earlier methods, such as Sanger sequencing, Northern blotting, and qRT-PCR, were both labor and time-intensive and could only analyze a small portion of the transcriptome. The introduction of high-throughput methods, such as RNA sequencing and its variants, have transformed transcriptomic research (Daines et al., 2011; Tang et al., 2009; Wang et al., 2009). RNA sequencing now allows for comprehensive profiling of gene expression, providing a broad and detailed view of transcriptomic dynamics. These advancements have significantly improved our capacity to study complex biological processes and diseases at a molecular level, advancing our understanding of various complex conditions.

Ribosomal RNA (rRNA), the most abundant type of RNA, typically constitutes about 80% of total RNA (Singer & Berg, 1991), posing a significant challenge for profiling the RNA types relevant for gene expression studies. Therefore, it must be removed from total RNA before sequencing, to allow for the enrichment of high-value targets (Huang et al., 2011; O’Neil et al., 2013). Currently, two methods exist for rRNA removal: polyA-enrichment and rRNA-depletion. PolyA-enrichment is widely used because of its cost-effectiveness and ability to work effectively with low sequencing depth, often providing better coverage compared to rRNA depletion at equivalent sequencing depth. Unfortunately, polyA-enrichment sometimes leads to a 3’ end bias (Tariq et al., 2011) which renders it less effective for low-quality RNA or Formalin-Fixed Paraffin-Embedded (FFPE) tissue samples, which generally have degraded RNA (Kellman et al., 2021; W. Zhao et al., 2014). While this means that rRNA that lack polyA tail are eliminated, it also leads to the loss of many non-coding RNA (ncRNA) lacking a polyA tail. rRNA-depletion, which focuses on the elimination of rRNA rather than the selection of a particular class of RNA serves as a much better alternative for low-quality and degraded RNA while also enriching the non-coding RNA that would otherwise be lost in the polyA-selection method (Cui et al., 2010; Kissopoulou et al., 2013). Depletion of rRNA generally depends on the principle of hybridization of single-stranded rDNA probes to rRNA. Thus, for efficient rRNA depletion and minimal off-target activity, these probes need to be highly specific for the species in which rRNA depletion must be done. This restricts the use of the kits to the intended organisms and those that share their conserved rRNA regions (Kraus et al., 2019). rRNA sequences are relatively conserved across eukaryotes, especially higher eukaryotes. Hence, kits developed for humans also work for rats and mice. However, in the case of insects, the ribosome biogenesis pathways are different. The 28S rDNA has 2 types of insertions; while Type I insertion resembles mammalian rDNA with a single block for 28S rDNA, Type II insertion has 2 blocks of 28S rRNA that are separated from each other by 5.4 kb of DNA (Glover & Hogness, 1977; Pellegrini et al., 1977; Tautz et al., 1988; White & Hogness, 1977). 28S rRNA is transcribed from these rDNA along with 5.8S and 18S as a single large precursor. Following splicing, the 28S rRNA undergoes further processing through an unknown mechanism which leads to fragmentation of the 28S rRNA into α and ß fragments (Dawid & Wellauer, 1978; Kidd & Glover, 1981; Winnebeck et al., 2010). Due to this fragmentation of the insect rRNA during processing, the commonly available kits used for vertebrate rRNA depletion, such as RiboZero and RiboZero Gold do not completely remove 5S rRNA or 28S rRNA, which leads to low mapping and sequence coverage. Thus, most commercially available kits do not recommend using kits developed for a particular organism or group of organisms to be used for rRNA depletion in other organisms (Figure 1A). This inefficiency can negatively impact downstream analyses, such as sequence mapping and coverage. Due to the lack of an efficient protocol, several previous studies have performed rRNA depletion using commercial kits designed for vertebrates in *Drosophila* (Table 1).

**Figure 1:**
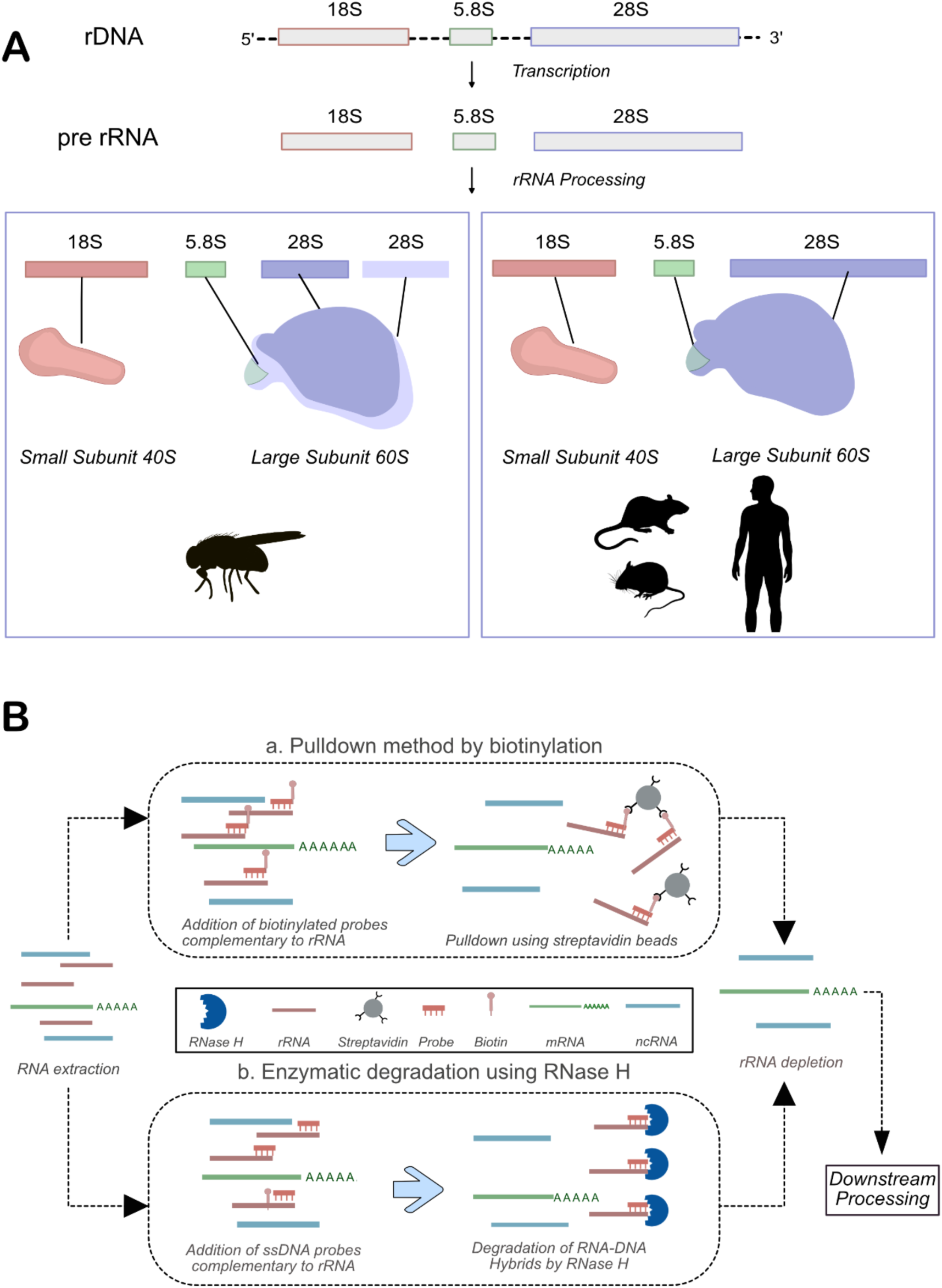
rRNA biogenesis and depletion methods. (A) Highlighting key differences in rRNA biogenesis pathways in Drosophila and Mammals. While mammals have one fragment of 28S rRNA, Drosophila 28S rRNA has two fragments: α and ß. (B) A schematic for using biotinylated probes or RNase H for rRNA-depletion.

**Table 1:** rRNA-depletion kits previously used for Drosophila samples.

| Name of Kit | Recommended Species | Notes | Reference |
| --- | --- | --- | --- |
| Tecan Ovation® RNA-Seq System V2 | - | cDNA synthesis without polyA selection or rRNA depletion | (Hughes et al., 2012) |
| Zymo-Seq RiboFree Total RNA library Kit | Any | 20% of Insect rRNA remains undepleted | (Talross & Carlson, 2023) |
| Illumina Ribo-Zero™ Gold Kit | HMR | Currently Unavailable | (M.-J. M. Chen et al., 2016) |
| Illumina Ribo-Zero rRNA removal kit | HMR | Currently Unavailable | (Pritykin et al., 2017) |
| Thermo Fisher Scientific RiboMinus™ Eukaryote Kit for RNA-Seq | HMR | Low compatibility with *Drosophila* as stated by the manufacturer | (Lefebvre et al., 2017; Lim et al., 2013; Mugat et al., 2015) |

(HMR – Human, Mouse and Rat)

However, due to these reasons mentioned earlier, kits developed for mammals that were used in earlier studies are no longer available or not recommended for use with *Drosophila* samples. However, there are a few promising developments in *Drosophila*-specific rRNA depletion kits like Zymo’s Seq RiboFree Total RNA Library Kit, which claims to remove approximately 80% of *Drosophila* rRNA. Also, Qiagen has developed a QIAseq FastSelect –rRNA Fly Kit specifically for *Drosophila* rRNA removal. Despite the availability and time-saving benefits of commercial kits, they are often inefficient and exorbitantly priced, leading some labs to opt for in-house preparation of rRNA-depletion methods.

There are two primary methods for removing rRNA, each with its own set of advantages and limitations. The first approach, known as pulldown, involves the use of biotinylated DNA oligos that are complementary to rRNA sequences. These oligos hybridize with the rRNA, and the resulting RNA-DNA hybrids are captured using streptavidin-conjugated beads. Once bound, the rRNA can be efficiently removed (Bhagwat et al., 2014; Z. Chen & Duan, 2011; Culviner et al., 2020). While this method is highly specific and effective for rRNA depletion, it is less suitable for fragmented or degraded RNA samples. The high specificity of the pulldown probes means that if the RNA is fragmented, only portions of the rRNA may bind to the probes, leaving residual rRNA in the sample. Additionally, it requires a large initial amount of starting material, which may not always be feasible. An alternative method is enzymatic depletion, which also relies on the formation of RNA-DNA hybrids (Baldwin et al., 2021). In this case, the RNase H enzyme is employed to degrade the rRNA specifically bound to the complementary DNA oligos (Figure 1B). This method has the advantage of being effective even with low-quality or degraded RNA (Wahl et al., 2022). However, different species require the design of species-specific probes for optimal performance. Both techniques have been used to develop in-house rRNA-depletion kits for *Drosophila*. Biotinylated probes have been developed to remove various rRNA fractions ranging from 2S to 28S rRNA (Fowler et al., 2018; Thompson et al., 2020). RNase H dependent enzymatic degradation has also been used for the depletion of 5.8S, 18S and 28S rRNA (Haugen et al., 2024). However, the probe sequences have not been made publicly available.

To address these challenges, we have designed rRNA probes specific for *Drosophila melanogaster* rRNA sequences and demonstrate effective and reproducible depletion of rRNA from adult *Drosophila* brain samples. We make the probe sequences freely available for use by the research community. Our study provides a comprehensive protocol for rRNA-depletion using the RNase H method, from probe design to data analysis, enabling other researchers to replicate and adapt this approach for their studies in *Drosophila melanogaster* and other closely related species.

## METHODS

### Fly Rearing and Genetics

*Drosophila* Canton-S stocks were maintained at 25 ±1°C on cornmeal-sugar-agar media.

### RNA Extraction

5-day-old adult *Drosophila* adult brains (N=15) were dissected in 1X PBS and RNA was extracted using TRIzol (Thermo Fisher Scientific) following the manufacturer’s protocol. The quality of RNA was assessed using Agilent Tapestation 4200 before proceeding with cDNA library preparation and sequencing.

### Probe Design and reconstitution

Sequences for 5S (URS00003B4856_7227), 5.8S (URS00005FF212_7227), 18S (URS0000A575BB_7227) and 28S (URS0000A53741_7227) rRNA were sourced from RNAcentral (Sweeney et al., 2019). Probes were designed using the NEBNext Custom RNA Depletion Design Tool. These probes were analyzed using BLAST to assess potential off-target effects, employing default parameters with specific adjustments for “Organism” (*Drosophila melanogaster*) and “Optimize for” (blastn) (Altschul et al., 1990). Subsequently, a final list of probes to cover the entire set of rDNA (Table S1) was synthesized by Integrated DNA Technologies, Pvt. Ltd. and reconstituted in nuclease-free water to achieve a stock concentration of 250 µM. These probes were then pooled together, resulting in a final concentration of 2 µM per probe.

### Depletion of rRNA using custom probes

For Poly(A)-enrichment of mRNA, Illumina libraries were prepared using NEBNext Ultra II Directional RNA Library Prep kit (E7765L) and sequenced with Illumina NovaSeq 6000 system. rRNA-depletion was performed as recommended by NEBNext RNA Depletion Core Reagent protocol. Briefly, the RNA integrity was confirmed on an Agilent Tapestation 4200 and samples with at least 7.0 RIN values were used for experiments. The total RNA (500-1000 ng), 2 μL of the 2μM probe pool, 2 μL of the 2μM NEBNext probe hybridization buffer and nuclease-free water were mixed to a total volume of 15 μL. This mixture was then incubated at 95°C for 2 min, a touchdown from 95°C to 22°C (0.1°C/sec) and a 5-min hold at 22°C. We then set up a reaction with 2 μL NEBNext RNase H Reaction Buffer, 2 μL NEBNext Thermostable RNase H and 1 μL nuclease-free water and incubated for 30 min at 50°C. The RNase H treated sample was digested with DNase I and purified using NEBNext RNA Sample Purification Beads as recommended by NEBNext RNA Depletion Core Reagent protocol. The concentrations of purified RNA samples were measured using the Qubit HS RNA Assay. The purified rRNA-depleted samples were taken for library preparation using NEBNext Ultra™ II Directional RNA Library Prep with Sample Purification Beads (Catalogue no-E7765L) as per the manufacturer’s protocol. The libraries were sequenced on the Illumina NovaSeq 6000 platform with a 2×100bp read length. Similarly, rRNA-depletion was performed as recommended by QIAseq FastSelect – rRNA Fly Kits protocol (Cat. No. / ID: 333262) using total RNA extracted from *Drosophila* S2 cells. The rRNA-depleted samples were processed using a similar protocol as described above.

### Data analysis

Sequenced files in FASTQ format were initially assessed for quality using FastQC (Andrews, 2017/2024). Trimmomatic was used to trim the reads as per the following parameters: *-phred33 HEADCROP:6 LEADING:25 TRAILING:25 AVGQUAL:25 MINLEN:19* (Bolger et al., 2014). Once high-quality reads were confirmed, alignment was performed against the *Drosophila* dm6 reference genome, using STAR (Dobin et al., 2013). The count matrix was generated using the featureCounts package (Liao et al., 2014). Differential expression analysis was conducted using the DESeq2 package in RStudio (Love et al., 2014). Visualization of results was achieved with EnhancedVolcano for volcano plots and pheatmap for heatmaps (Raivo Kolde, 2010). Picard (*Broadinstitute/Picard*, 2014/2024) was utilized to calculate gene region coverage, while custom scripts were employed for generating Venn diagrams and bar plots. Ribodetector was used to estimate the rRNA content in each sample (Deng et al., 2022, p. 2). The softwares used in the study are listed in Table S4.

## RESULTS

### Design and selection of custom probes for *Drosophila* rRNA-depletion

The effectiveness of rRNA removal using RNase H relies on successful hybridization between rRNA and single-stranded DNA probes with sequence homology. Poorly designed probes can impact downstream steps and compromise data quality, making probe design a critical step in rRNA depletion. For this purpose, we downloaded the sequences of *Drosophila melanogaster* 5S (URS00003B4856_7227), 5.8S (URS00005FF212_7227), 18S (URS0000A575BB_7227) and 28S (URS0000A53741_7227) rRNA from RNACentral and designed specific probes using the NEBNext Custom RNA Depletion Design Tool. This process generated 2, 2, 34, and 64 probes for 5S, 5.8S, 18S, and 28S rRNA, respectively. Since RNase H degrades RNA-DNA hybrid regardless of sequence, non-specific probes could inadvertently deplete sequences within our transcriptome of interest. To prevent this, we performed sequence homology searches using BLAST for all the probe sequences, ensuring that non-target RNA with homology to our probes would not be removed unintentionally.

Using the NEBNext Custom RNA Depletion Design Tool, our initial design generated 164 probes, many of which had overlapping sequences. After manual curation of the overlapping probes, we finalized a set of 102 probes (Table S1) that comprehensively covered the entire lengths of the 5S, 5.8S, 18S, and 28S rRNAs with minimal gaps.

### Custom-generated probes for RNase H based rRNA-depletion performed better than the commercial kit

RNA was extracted from *Drosophila* brains, and its quality was evaluated using an Agilent Tapestation 4200. The freshly isolated total RNA showed a prominent rRNA peak corresponding to the 18S size (Figure 2B-i). Unlike mammalian rRNA, which produces two distinct peaks at approximately 1.9 kb and 5 kb for 18S and 28S respectively, *Drosophila* 28S rRNA is naturally fragmented into 28*S* α and 28S ß which migrates close to the 18S rRNA, resulting in two prominent peaks at approximately 2 kb (Winnebeck et al., 2010). This total RNA was used for rRNA-depletion using the synthesized custom rRNA probes and RNase H as described in the methods section. The rRNA-depleted samples were evaluated using an Agilent Tapestation 4200, which showed a complete absence of peaks corresponding to 18S and 28S rRNA bands post-depletion (Figure 2B-ii). An equivalent quality of total RNA extracted from S2 cells was used to deplete rRNA using the QIAseq FastSelect –rRNA Fly Kit. In parallel, we also enriched polyA RNA from both samples to compare the levels of rRNA and ncRNA with respect to rRNA-depleted samples.

**Figure 2:**
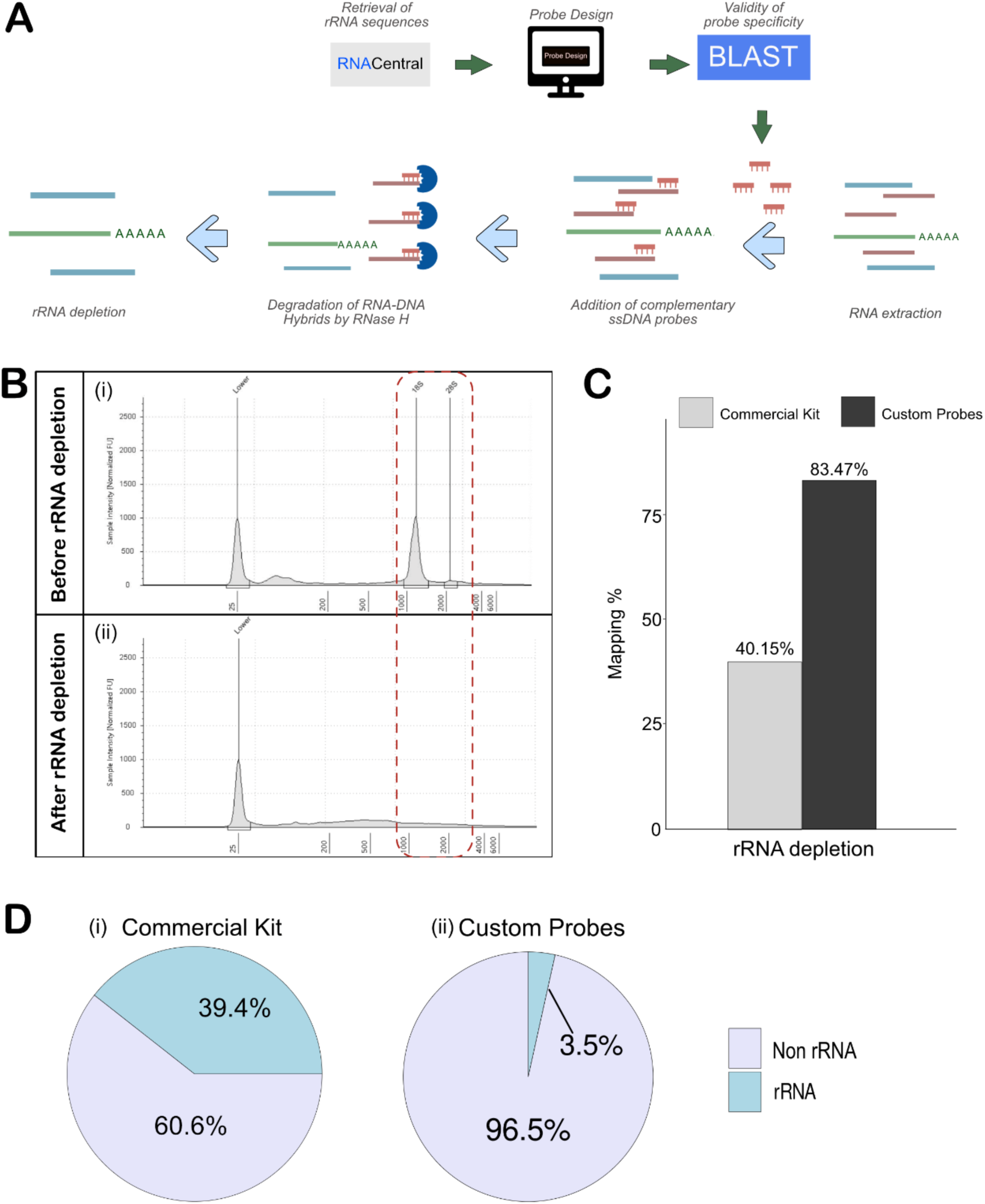
Efficient depletion of rRNA with RNase H using custom-generated probes. (A) Schematic workflow for probe design and rRNA depletion using RNase H. (B) Tape station analysis shows the depletion of 18S and 28S rRNAs. (C) Mapping percentages of samples sequenced using the RNase H method and Qiaseq FastSelect - fly rRNA-depletion kit. (D) Mapped rRNA read percentage from the RNase H method and Qiaseq Fastselect - fly rRNA-depletion kit.

Next-generation sequencing (NGS) libraries were generated for rRNA-depleted and polyA-selected RNA using NEBNext Ultra™ II Directional RNA Library Prep Kit. The libraries were sequenced on Illumina NovaSeq 6000, and the reads and quality scores are tabulated in Table S2. FastQC was used to assess the quality of sequenced reads that consistently returned a high quality across all samples, with per-base sequence quality scores exceeding 35.

With the data quality confirmed, the sequencing reads were trimmed as described in the methods section, before mapping to the *Drosophila melanogaster* reference genome (dm6) using STAR. In the samples where custom probes were used for rRNA depletion, the mapping was about 83.47%, which was quite comparable to the polyA-enriched sample. However, the commercial kit-based rRNA depletion removed the rRNA inefficiently, leading to just ∼ 40% mapping as compared to the 83.2% mapping in the polyA-enriched sample (Figure 2C, Table S2). The effectiveness of rRNA removal was determined using Ribodetector, which showed that samples prepared with the QIAseq FastSelect –rRNA Fly kit retained ∼39% rRNA, which explains the poor mapping rate (Figure 2D-i). On the other hand, the custom probe-based method retained just 3.5%, showing efficient rRNA depletion (Figure 2D-ii). Given the lack of a sufficient number of publications using the QIAseq FastSelect -rRNA Fly kit, it might need optimization. However, our findings indicate that, in its current iteration, the probe-based RNase H method provides superior performance for rRNA depletion.

### Non-coding RNAs were identified in the rRNA-depleted samples

The purpose of using rRNA depletion sequencing rather than polyA-selection is to capture a broader population of ncRNAs, which typically lack polyA tail. To evaluate whether the rRNA-depletion method successfully enriched the non-coding transcriptome, we generated count matrices for both rRNA-depleted and polyA-enriched datasets from BAM files using featureCounts. We then performed differential expression analysis by comparing the rRNA count matrix to the polyA count matrix. A volcano plot illustrated the enrichment of RNAs in the rRNA-depleted samples, with several ncRNAs such as CR434459, snRNA:7SK, and hsr-omega prominently upregulated in rRNA-depleted samples (Figure 3A). A list of these upregulated genes, including their log2 fold changes and adjusted p-values, can be found in Table S3. A heatmap to display the differences in enriched genes between the rRNA-depleted and polyA-selection samples is shown in Figure 3B.

**Figure 3:**
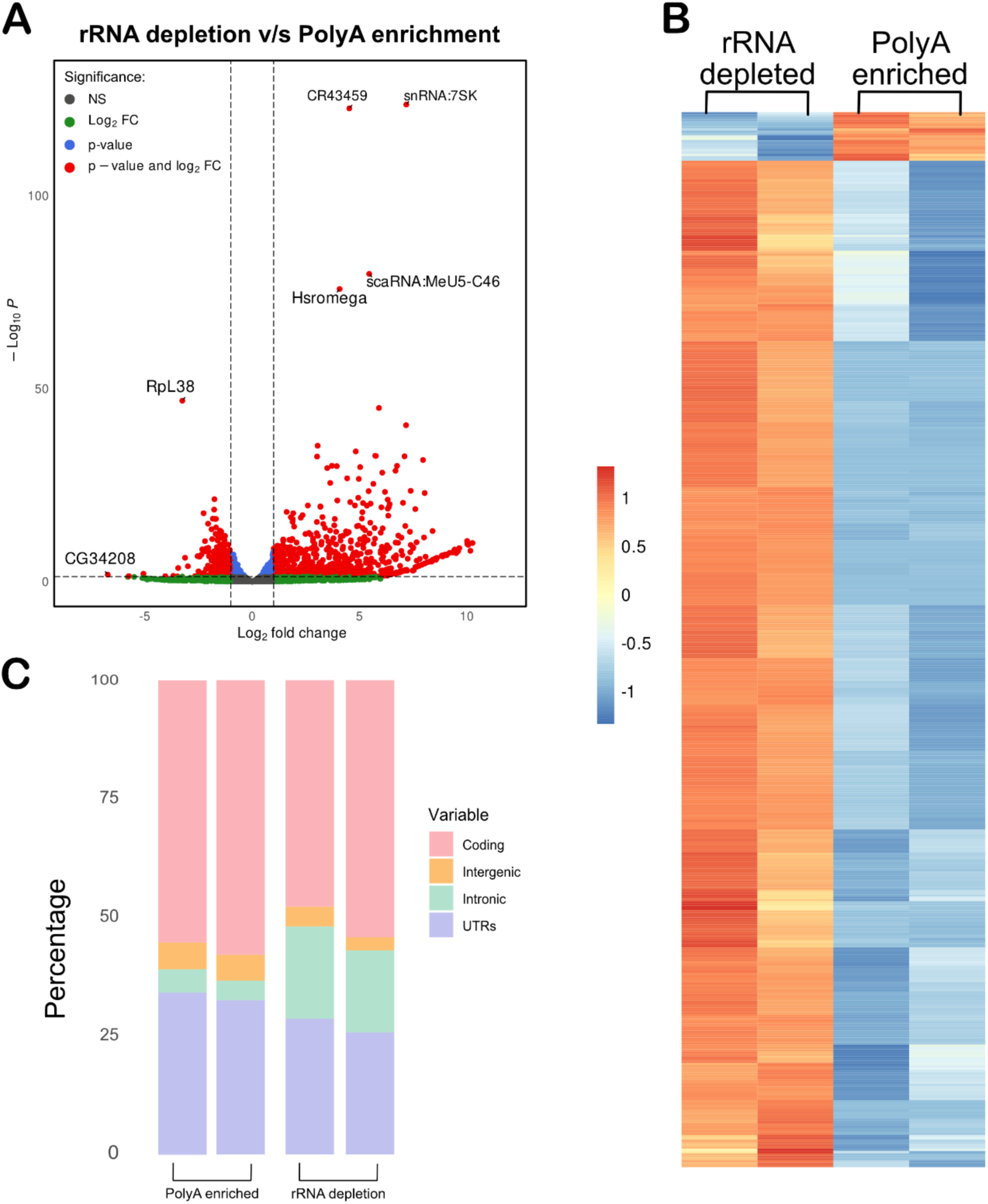
Non-coding RNAs were enriched in the rRNA-depleted samples. (A) A Volcano plot showing ncRNA enriched in rRNA-depleted samples as compared to the polyA-enriched samples. (B) Heat map of genes with >2-fold change and <0.05 padj value (Fold change values in Table S3). (C) rRNA-depletion increased intronic coverage as compared to the polyA-enriched samples.

Furthermore, we classified the top 15 ncRNA, ranking them by log2 fold change while excluding those with adjusted p-values greater than 0.05. Most of the identified ncRNA were long non-coding RNAs (lncRNAs), such as CR44157 and CR44474, followed by small nucleolar RNAs (snoRNAs) (Table 2).

**Table 2:** Top 15 non-coding RNA.

| Name | Classification |
| --- | --- |
| snoRNA: CG32479-b | sno RNA |
| snoRNA: Or-aca4 | sno RNA |
| mir-4969 | miRNA |
| mir-2279 | miRNA |
| CR44157 | lncRNA |
| mir-317 | miRNA |
| CR44474 | lncRNA |
| CR43399 | lncRNA |
| CR45256 | lncRNA |
| snoRNA: Psi28S-2622 | sno RNA |
| CR44279 | lncRNA |
| mir-4982 | miRNA |
| CR44496 | lncRNA |
| CR44034 | lncRNA |
| CR44070 | lncRNA |

In addition to the detection of mRNA and ncRNA, rRNA depletion can capture nascent transcripts, and thus, there is a possibility of higher intronic sequences in rRNA-depletion as compared to polyA-enrichment samples (S. Zhao et al., 2018). To confirm this, we used Picard to assess coverage across different genomic regions in both rRNA depleted and polyA samples. Intronic sequences were notably more abundant in the rRNA depleted samples compared to the polyA samples (Figure 3C). This observation suggests that, while not highly efficient, the rRNA depletion method can detect nascent mRNAs.

Given this, rRNA depletion can be considered a suitable approach for most studies, particularly considering the increasing recognition of the functional importance of ncRNAs. Unless the study specifically requires the detection of low-abundance mRNA targets, rRNA depletion offers a compelling alternative, enabling the exploration of both coding and non-coding RNA dynamics.

## DISCUSSION

Despite being a vital model organism, *Drosophila* has been largely overlooked when it comes to the development of rRNA-depletion kits. Most commercially available kits are specifically optimised for mammalian systems, primarily humans, mice, and rats (HMR). This bias has forced researchers working with *Drosophila* to adapt HMR-optimised kits for rRNA depletion in their studies. However, this approach is suboptimal due to significant differences between *Drosophila* and HMR rRNA structures, particularly the fragmented nature of the 28S rRNA in *Drosophila* compared to the single, continuous 28S rRNA found in mammals. These differences affect the efficiency of rRNA depletion in *Drosophila*, highlighting the need for a species-specific solution.

To fill this gap, we developed a *Drosophila*-specific rRNA depletion method. Although some laboratories have created rRNA depletion techniques for *Drosophila*, their complete probe sequences are unavailable, and their efficiency has not been thoroughly validated or compared to commercial kits (Haugen et al., 2024; Thompson et al., 2020). Existing commercial kits often rely on biotinylated probes, which are inefficient for fragmented or low-quality RNA (Culviner et al., 2020). To address these challenges, we designed custom probes specifically targeting *Drosophila* rRNA and paired them with RNase H for efficient depletion. This approach is optimized to enhance efficiency and produces rRNA-depleted RNA suitable for diverse downstream applications, including qRT-PCR, microarray analysis, and NGS.

The NEB Probe Designer tool, used for synthesizing rRNA probes, generates multiple probes with overlapping sequences. To minimize costs and avoid redundancy, these overlapping probes were manually reviewed and eliminated. Due to the short sequence of 2S rRNA (∼30 nucleotides), the tool did not design a probe for it. However, we did not observe 2S rRNA in our sequencing reads, as it might have been depleted in the RNA purification step following enzymatic digestions.

Further, this method is best suited for the identification of ncRNA in fly tissues. Differential analysis identified several ncRNAs, particularly lncRNAs, were significantly enriched in the rRNA-depleted samples compared to the polyA-enriched ones (Figure 3A). Previous studies have reported that nascent mRNAs, which typically lack a polyA tail, are not efficiently detected using polyA-enriched samples (Yang et al., 2011; S. Zhao et al., 2018). However, rRNA depletion allows nascent RNA detection from the intronic reads (Figure 3C).

Interestingly, we also detected a few microRNAs (miRNAs) (Table 2). Given their small size (20–22 nucleotides), miRNAs are typically not expected in rRNA-depleted samples and require specialized sequencing protocols that include specific size-selection steps for enrichment. We hypothesize that these detected miRNAs may belong to a subset of highly abundant ones.

Despite these strengths, a few limitations of the method should be noted. Samples sequenced following rRNA depletion and mapping to the reference genome had a small percentage of reads that were too short to align effectively (”% of reads unmapped: too short”). These could be due to leftover rRNA fragments from depletion. Further STAR documentation does note that “poor sequencing quality, over-trimming, or contamination with ribosomal RNA fragments can yield reads classified as *too short to map*”. Despite this, we obtained mapping percentages of custom probes mediated rRNA depleted samples that were comparable to the polyA selection method (Figure 2C).

These findings underscore the suitability of rRNA depletion for capturing both coding and non-coding RNA populations, including nascent RNA. Given the high-quality results observed, this method is a robust alternative to commercial kits, and we are confident that this custom probe-based rRNA depletion method will be used more often while sequencing *Drosophila* species as well as closely related insect transcriptomes.

## Supporting information

Table S1: List of custom probes used for RNase H mediated rRNA-depletion

Table S2: NGS data mapping details

Table S3: Differential Expression analysis results

Table S4: List of software used in the study

## EXTENDED DATA

Table S1: List of custom probes used for RNase H mediated rRNA-depletion

Table S2: NGS data mapping details

Table S3: Differential Expression analysis results

Table S4: List of software used in the study

## DATA AVAILABILITY

The NGS data used in the study can be accessed at GEO ID: GSE282990

## ACKNOWLEDGMENTS

We thank members of the Bakthavachalu lab and Amanjot Singh, for their useful comments on the manuscript. The fly facility at Bangalore Life Science Cluster (BLiSC) provided support with fly stock supply as well as generation of transgenics and next-generation genomics facility at BLiSC provided NGS service. Finally, we would like to thank the PARAM Himalaya cluster at IIT Mandi for providing the computational resources and technical support for carrying out various analysis.

## AUTHOR CONTRIBUTIONS

Conceptualization: BB

Methodology: OK, ST, AC, AP and BB

Investigation: OK, ST, AC and AP

Funding Acquisition: BB

Project Administration: BB

Supervision: BB

Writing – original draft: OK and BB

Writing – review and editing: ST, AP and BB

## REFERENCES

1. Adams, M. D., Celniker, S. E., Holt, R. A., Evans, C. A., Gocayne, J. D., Amanatides, P. G., Scherer, S. E., Li, P. W., Hoskins, R. A., Galle, R. F., George, R. A., Lewis, S. E., Richards, S., Ashburner, M., Henderson, S. N., Sutton, G. G., Wortman, J. R., Yandell, M. D., Zhang, Q., … Venter, J. C. (2000). The Genome Sequence of Drosophila melanogaster. Science, 287(5461), 2185–2195. 10.1126/science.287.5461.2185

2. Altschul, S. F., Gish, W., Miller, W., Myers, E. W., & Lipman, D. J. (1990). Basic local alignment search tool. Journal of Molecular Biology, 215(3), 403–410. 10.1016/S0022-2836(05)80360-2

3. Andrews, S. (2024). *S-andrews/FastQC* [Java]. https://github.com/s-andrews/FastQC (Original work published 2017)

4. Baldwin, A., Morris, A. R., & Mukherjee, N. (2021). An Easy, Cost-Effective, and Scalable Method to Deplete Human Ribosomal RNA for RNA-seq. Current Protocols, 1(6), e176. 10.1002/cpz1.176

5. Banfi, S., Borsani, G., Rossi, E., Bernard, L., Guffanti, A., Rubboli, F., Marchitiello, A., Giglio, S., Coluccia, E., Zollo, M., Zuffardi, O., & Ballabio, A. (1996). Identification and mapping of human cDNAs homologous to Drosophila mutant genes through EST database searching. Nature Genetics, 13(2), 167–174. 10.1038/ng0696-167

6. Bhagwat, A. A., Ying, Z. I., & Smith, A. (2014). Evaluation of Ribosomal RNA Removal Protocols for *Salmonella* RNA-Seq Projects. Advances in Microbiology, 2014. 10.4236/aim.2014.41006

7. Bolger, A. M., Lohse, M., & Usadel, B. (2014). Trimmomatic: A flexible trimmer for Illumina sequence data. Bioinformatics, 30(15), 2114–2120. 10.1093/bioinformatics/btu170

8. Bradley, R. K., & Anczuków, O. (2023). RNA splicing dysregulation and the hallmarks of cancer. Nature Reviews Cancer, 23(3), 135–155. 10.1038/s41568-022-00541-7

9. *Broadinstitute/picard*. (2024). [Java]. Broad Institute. https://github.com/broadinstitute/picard (Original work published 2014)

10. Chen, M.-J. M., Chen, L.-K., Lai, Y.-S., Lin, Y.-Y., Wu, D.-C., Tung, Y.-A., Liu, K.-Y., Shih, H.-T., Chen, Y.-J., Lin, Y.-L., Ma, L.-T., Huang, J.-L., Wu, P.-C., Hong, M.-Y., Chu, F.-H., Wu, J.-T., Li, W.-H., & Chen, C.-Y. (2016). Integrating RNA-seq and ChIP-seq data to characterize long non-coding RNAs in Drosophila melanogaster. BMC Genomics, 17(1), 220. 10.1186/s12864-016-2457-0

11. Chen, Z., & Duan, X. (2011). Ribosomal RNA Depletion for Massively Parallel Bacterial RNA-Sequencing Applications. In Y. M. Kwon & S. C. Ricke (Eds.), High-Throughput Next Generation Sequencing: Methods and Applications (pp. 93–103). Humana Press. 10.1007/978-1-61779-089-8_7

12. Cui, P., Lin, Q., Ding, F., Xin, C., Gong, W., Zhang, L., Geng, J., Zhang, B., Yu, X., Yang, J., Hu, S., & Yu, J. (2010). A comparison between ribo-minus RNA-sequencing and polyA-selected RNA-sequencing. Genomics, 96(5), 259–265. 10.1016/j.ygeno.2010.07.010

13. Culviner, P. H., Guegler, C. K., & Laub, M. T. (2020). A Simple, Cost-Effective, and Robust Method for rRNA Depletion in RNA-Sequencing Studies. mBio, 11(2), 10.1128/mbio.00010-20.

14. Daines, B., Wang, H., Wang, L., Li, Y., Han, Y., Emmert, D., Gelbart, W., Wang, X., Li, W., Gibbs, R., & Chen, R. (2011). The Drosophila melanogaster transcriptome by paired-end RNA sequencing. Genome Research, 21(2), 315–324. 10.1101/GR.107854.110

15. Dawid, I. B., & Wellauer, P. K. (1978). Ribosomal DNA and Related Sequences in Drosophila melanogaster. Cold Spring Harbor Symposia on Quantitative Biology, 42, 1185–1194. 10.1101/SQB.1978.042.01.119

16. Deng, Z.-L., Münch, P. C., Mreches, R., & McHardy, A. C. (2022). Rapid and accurate identification of ribosomal RNA sequences via deep learning. Nucleic Acids Research, 50(10), e60. 10.1093/nar/gkac112

17. Dobin, A., Davis, C. A., Schlesinger, F., Drenkow, J., Zaleski, C., Jha, S., Batut, P., Chaisson, M., & Gingeras, T. R. (2013). STAR: Ultrafast universal RNA-seq aligner. Bioinformatics, 29(1), 15–21. 10.1093/bioinformatics/bts635

18. Fortini, M. E., Skupski, M. P., Boguski, M. S., & Hariharan, I. K. (2000). A Survey of Human Disease Gene Counterparts in the Drosophila Genome. Journal of Cell Biology, 150(2), F23–F30. 10.1083/jcb.150.2.F23

19. Fowler, E. K., Mohorianu, I., Smith, D. T., Dalmay, T., & Chapman, T. (2018). Small RNA populations revealed by blocking rRNA fragments in Drosophila melanogaster reproductive tissues. PLOS ONE, 13(2), e0191966. 10.1371/journal.pone.0191966

20. Glover, D. M., & Hogness, D. S. (1977). A novel arrangement of the 18S and 28S sequences in a repeating unit of drosophila melanogaster rDNA. Cell, 10(2), 167–176. 10.1016/0092-8674(77)90212-4

21. Goodman, L. D., & Bellen, H. J. (2022). Recent insights into the role of glia and oxidative stress in Alzheimer’s disease gained from *Drosophila*. Current Opinion in Neurobiology, 72, 32–38. 10.1016/j.conb.2021.07.012

22. Hardin, P. E., Hall, J. C., & Rosbash, M. (1990). Feedback of the Drosophila period gene product on circadian cycling of its messenger RNA levels. Nature, 343(6258), 536–540. 10.1038/343536a0

23. Harvey, K. F., Pfleger, C. M., & Hariharan, I. K. (2003). The Drosophila Mst Ortholog, hippo, Restricts Growth and Cell Proliferation and Promotes Apoptosis. Cell, 114(4), 457–467. 10.1016/S0092-8674(03)00557-9

24. Haugen, R. J., Barnier, C., Elrod, N. D., Hua, L. U. O., Jensen, M. K., Ping, J. I., Smibert, C. A., Lipshitz, H. D., Wagner, E. J., Freddolino, P. L., & Goldstrohm, A. C. (2024). Regulation of the Drosophila transcriptome by Pumilio and the CCR4–NOT deadenylase complex. RNA, 30(7), 866–890. 10.1261/RNA.079813.123

25. Huang, R., Jaritz, M., Guenzl, P., Vlatkovic, I., Sommer, A., Tamir, I. M., Marks, H., Klampfl, T., Kralovics, R., Stunnenberg, H. G., Barlow, D. P., & Pauler, F. M. (2011). An RNA-Seq Strategy to Detect the Complete Coding and Non-Coding Transcriptome Including Full-Length Imprinted Macro ncRNAs. PLOS ONE, 6(11), e27288. 10.1371/journal.pone.0027288

26. Hughes, M. E., Grant, G. R., Paquin, C., Qian, J., & Nitabach, M. N. (2012). Deep sequencing the circadian and diurnal transcriptome of Drosophila brain. Genome Research, 22(7), 1266–1281. 10.1101/gr.128876.111

27. Kellman, B. P., Baghdassarian, H. M., Pramparo, T., Shamie, I., Gazestani, V., Begzati, A., Li, S., Nalabolu, S., Murray, S., Lopez, L., Pierce, K., Courchesne, E., & Lewis, N. E. (2021). Multiple freeze-thaw cycles lead to a loss of consistency in poly(A)-enriched RNA sequencing. BMC Genomics, 22(1), 69. 10.1186/s12864-021-07381-z

28. Kidd, S. J., & Glover, D. M. (1981). *Drosophila melanogaster* ribosomal DNA containing type II insertions is variably transcribed in different strains and tissues. Journal of Molecular Biology, 151(4), 645–662. 10.1016/0022-2836(81)90428-9

29. Kim, S. K., Tsao, D. D., Suh, G. S. B., & Miguel-Aliaga, I. (2021). Discovering signaling mechanisms governing metabolism and metabolic diseases with Drosophila. Cell Metabolism, 33(7), 1279–1292. 10.1016/j.cmet.2021.05.018

30. Kissopoulou, A., Jonasson, J., Lindahl, T. L., & Osman, A. (2013). Next Generation Sequencing Analysis of Human Platelet PolyA+ mRNAs and rRNA-Depleted Total RNA. PLOS ONE, 8(12), e81809. 10.1371/journal.pone.0081809

31. Kraus, A. J., Brink, B. G., & Siegel, T. N. (2019). Efficient and specific oligo-based depletion of rRNA. Scientific Reports, 9(1), 12281. 10.1038/s41598-019-48692-2

32. Lefebvre, F. A., Benoit Bouvrette, L. P., Bergalet, J., & Lécuyer, E. (2017). Biochemical Fractionation of Time-Resolved *Drosophila* Embryos Reveals Similar Transcriptomic Alterations in Replication Checkpoint and Histone mRNA Processing Mutants. Journal of Molecular Biology, 429(21), 3264–3279. 10.1016/j.jmb.2017.01.022

33. Liao, Y., Smyth, G. K., & Shi, W. (2014). featureCounts: An efficient general purpose program for assigning sequence reads to genomic features. Bioinformatics, 30(7), 923–930. 10.1093/bioinformatics/btt656

34. Lim, S. J., Boyle, P. J., Chinen, M., Dale, R. K., & Lei, E. P. (2013). Genome-wide localization of exosome components to active promoters and chromatin insulators in Drosophila. Nucleic Acids Research, 41(5), 2963–2980. 10.1093/nar/gkt037

35. Link, N., & Bellen, H. J. (2020). Using Drosophila to drive the diagnosis and understand the mechanisms of rare human diseases. Development, 147(21), dev191411. 10.1242/dev.191411

36. Love, M. I., Huber, W., & Anders, S. (2014). Moderated estimation of fold change and dispersion for RNA-seq data with DESeq2. Genome Biol., 15(12). 10.1186/s13059-014-0550-8

37. Mirzoyan, Z., Sollazzo, M., Allocca, M., Valenza, A. M., Grifoni, D., & Bellosta, P. (2019). Drosophila melanogaster: A model organism to study cancer. Frontiers in Genetics, 10, 425859. 10.3389/FGENE.2019.00051/BIBTEX

38. Mugat, B., Akkouche, A., Serrano, V., Armenise, C., Li, B., Brun, C., Fulga, T. A., Vactor, D. V., Pélisson, A., & Chambeyron, S. (2015). MicroRNA-Dependent Transcriptional Silencing of Transposable Elements in Drosophila Follicle Cells. PLOS Genetics, 11(5), e1005194. 10.1371/journal.pgen.1005194

39. Nemeth, K., Bayraktar, R., Ferracin, M., & Calin, G. A. (2024). Non-coding RNAs in disease: From mechanisms to therapeutics. Nature Reviews Genetics, 25(3), 211–232. 10.1038/s41576-023-00662-1

40. Nüsslein-Volhard, C., & Wieschaus, E. (1980). Mutations affecting segment number and polarity in Drosophila. Nature, 287(5785), 795–801. 10.1038/287795a0

41. O’Neil, D., Glowatz, H., & Schlumpberger, M. (2013). Ribosomal RNA depletion for efficient use of RNA-seq capacity. *Current Protocols in Molecular Biology*, Chapter 4, Unit 4.19. 10.1002/0471142727.mb0419s103

42. Pellegrini, M., Manning, J., & Davidson, N. (1977). Sequence arrangement of the rDNA of Drosophila melanogaster. Cell, 10(2), 213–224. 10.1016/0092-8674(77)90215-X

43. Pritykin, Y., Brito, T., Schupbach, T., Singh, M., & Pane, A. (2017). Integrative analysis unveils new functions for the *Drosophila* Cutoff protein in noncoding RNA biogenesis and gene regulation. RNA, 23(7), 1097–1109. 10.1261/rna.058594.116

44. Raivo Kolde. (2010). *pheatmap: Pretty Heatmaps* (p. 1.0.12) [Dataset]. 10.32614/CRAN.package.pheatmap

45. Singer, Maxine., & Berg, P. (1991). Genes & genomes: A changing perspective. University Science Books.

46. Souidi, A., & Jagla, K. (2021). Drosophila Heart as a Model for Cardiac Development and Diseases. Cells, 10(11), Article 11. 10.3390/cells10113078

47. Sweeney, B. A., Petrov, A. I., Burkov, B., Finn, R. D., Bateman, A., Szymanski, M., Karlowski, W. M., Gorodkin, J., Seemann, S. E., Cannone, J. J., Gutell, R. R., Fey, P., Basu, S., Kay, S., Cochrane, G., Billis, K., Emmert, D., Marygold, S. J., Huntley, R. P., … Williams, K. P. (2019). RNAcentral: A hub of information for non-coding RNA sequences. Nucleic Acids Research, 47(D1), D221–D229. 10.1093/NAR/GKY1034

48. Talross, G. J. S., & Carlson, J. R. (2023). The rich non-coding RNA landscape of the Drosophila antenna. Cell Reports, 42(5). 10.1016/j.celrep.2023.112482

49. Tang, F., Barbacioru, C., Wang, Y., Nordman, E., Lee, C., Xu, N., Wang, X., Bodeau, J., Tuch, B. B., Siddiqui, A., Lao, K., & Surani, M. A. (2009). mRNA-Seq whole-transcriptome analysis of a single cell. Nature Methods, 6(5), 377–382. 10.1038/nmeth.1315

50. Tariq, M. A., Kim, H. J., Jejelowo, O., & Pourmand, N. (2011). Whole-transcriptome RNAseq analysis from minute amount of total RNA. Nucleic Acids Research, 39(18), e120. 10.1093/nar/gkr547

51. Tautz, D., Hancock, J. M., Webb, D. A., Tautz, C., & Dover, G. A. (1988). Complete sequences of the rRNA genes of Drosophila melanogaster. Molecular Biology and Evolution, 5(4), 366–376. 10.1093/oxfordjournals.molbev.a040500

52. Thompson, M. K., Kiourlappou, M., & Davis, I. (2020). Ribo-Pop: Simple, cost-effective, and widely applicable ribosomal RNA depletion. RNA, 26(11), 1731–1742. 10.1261/rna.076562.120

53. Tolwinski, N. S. (2017). Introduction: Drosophila—A Model System for Developmental Biology. Journal of Developmental Biology, 5(3). 10.3390/JDB5030009

54. Wahl, A., Huptas, C., & Neuhaus, K. (2022). Comparison of rRNA depletion methods for efficient bacterial mRNA sequencing. Scientific Reports, 12(1), 5765. 10.1038/s41598-022-09710-y

55. Wang, Z., Gerstein, M., & Snyder, M. (2009). RNA-Seq: A revolutionary tool for transcriptomics. Nature Reviews Genetics 2008 10:1, 10(1), 57–63. 10.1038/nrg2484

56. White, R. L., & Hogness, D. S. (1977). R loop mapping of the 18S and 28S sequences in the long and short repeating units of drosophila melanogaster rDNA. Cell, 10(2), 177–192. 10.1016/0092-8674(77)90213-6

57. Winnebeck, E. C., Millar, C. D., & Warman, G. R. (2010). Why does insect RNA look degraded? Journal of Insect Science, 10(1), 159. 10.1673/031.010.14119

58. Yang, L., Duff, M. O., Graveley, B. R., Carmichael, G. G., & Chen, L.-L. (2011). Genomewide characterization of non-polyadenylated RNAs. Genome Biology, 12(2), R16. 10.1186/gb-2011-12-2-r16

59. Zhao, S., Zhang, Y., Gamini, R., Zhang, B., & von Schack, D. (2018). Evaluation of two main RNA-seq approaches for gene quantification in clinical RNA sequencing: polyA+ selection versus rRNA depletion. Scientific Reports, 8(1), 4781. 10.1038/s41598-018-23226-4

60. Zhao, W., He, X., Hoadley, K. A., Parker, J. S., Hayes, D. N., & Perou, C. M. (2014). Comparison of RNA-Seq by poly (A) capture, ribosomal RNA depletion, and DNA microarray for expression profiling. BMC Genomics, 15(1), 419. 10.1186/1471-2164-15-419

